# Probing the causal role of prestimulus interregional synchrony for perceptual integration via tACS

**DOI:** 10.1101/044636

**Authors:** Rolandas Stonkus, Verena Braun, Jess Kerlin, Gregor Volberg, Simon Hanslmayr

## Abstract

The phase of prestimulus oscillations at 7-10 Hz has been shown to modulate perception of briefly presented visual stimuli. Specifically, a recent combined EEG-fMRI study suggested that a prestimulus oscillation at around 7 Hz represents open and closed windows for perceptual integration by modulating connectivity between lower order occipital and higher order parietal brain regions. We here utilized brief event-related transcranial alternating current stimulation (tACS) to specifically modulate this prestimulus 7 Hz oscillation, and the synchrony between parietal and occipital brain regions. To this end we tested for a causal role of this particular prestimulus oscillation for perceptual integration. The EEG was acquired at the same time allowing us to investigate frequency specific after effects phase-locked to stimulation offset. On a behavioural level our results suggest that the tACS did modulate perceptual integration, however, in an unexpected manner. On an electrophysiological level our results suggest that brief tACS does induce oscillatory entrainment, as visible in frequency specific activity phase-locked to stimulation offset. Together, our results do not strongly support a causal role of prestimulus 7 Hz oscillations for perceptual integration. However, our results suggest that brief tACS is capable of modulating oscillatory activity in a temporally sensitive manner.

## Introduction

Brain oscillations represent open and closed time windows for neural firing and thereby enable communication between distant neural populations^1^. In line with this hypothesis, a number of studies showed that the phase in the alpha/theta frequency band (7-10 Hz) at stimulus onset correlates with the likelihood of perceiving a briefly presented visual stimulus^2-7^. In other words, the chances of perceiving a visual stimulus in these studies closely followed a sine function depending on the phase of an ongoing oscillation at, or closely before, stimulus onset. In a recent EEG-fMRI study^4^ we replicated this well-documented relation between pre-stimulus phase and perception in a perceptual integration task (depicted in Figure 1). In that study we further demonstrated that interregional communication between lower visual processing areas in the left occipital cortex and higher order processing areas in the right intraparietal sulcus was modulated by pre-stimulus phase at 7 Hz, suggesting that neural communication between those two regions, and perceptual integration, is mediated by a 7 Hz oscillation. However, it is unclear to which extent this relationship between prestimulus inter-regional synchrony in the theta frequency band goes beyond correlation. In the current study we attempted to directly manipulate the level of prestimulus theta phase synchrony between the two regions (i.e. left occipital cortex and right parietal cortex) in a perceptual integration task. To this end we test for a causal relationship between prestimulus theta phase and perceptual integration.

**Figure 1.**
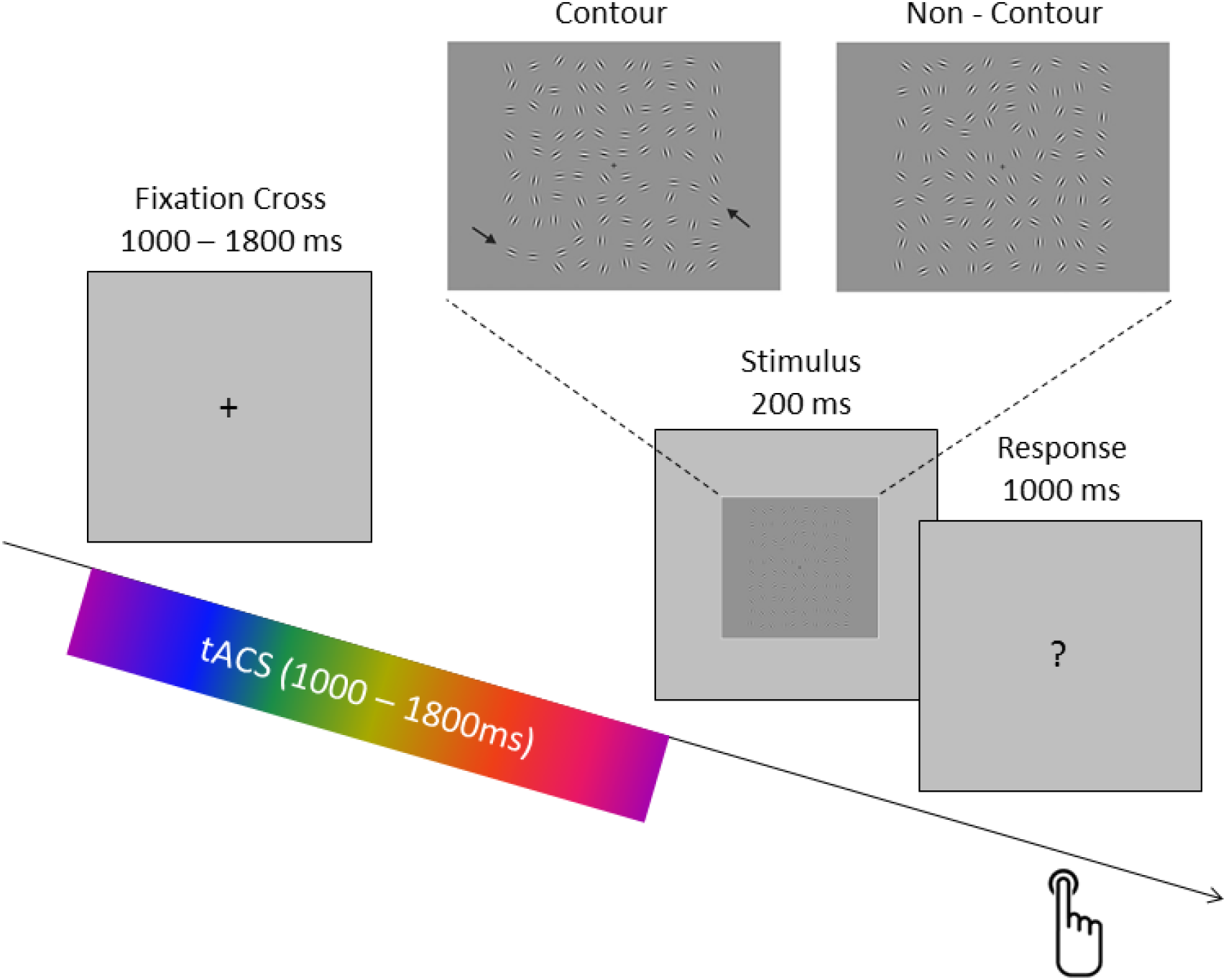
The perceptual integration task and the structure of a trial are depicted. The task of the subjects was to indicate via a button press whether a contour (upper left panel) or a non-contour stimulus (upper right panel) was shown (arrows for illustrative purposes only). Event-related tACS was delivered only during the variable interval presentation of a fixation cross. TACS conditions varied randomly between trials.

Transcranial alternating current stimulation (tACS) is a non-invasive electrical stimulation technique that injects relatively weak currents, oscillating at a particular frequency, via electrodes placed on the scalp^8,9^. A study in animals (i.e. ferrets) showed that the application of such weak alternating currents is indeed capable of entraining local network activity in the neocortex^10^. In the human brain, these results are supported by studies showing that behaviour can indeed be modulated by tACS in a way as would be predicted by entrainment^11-13^. Importantly, recent studies suggest that tACS is not only capable of influencing local oscillatory activity, but is also able to manipulate phase synchrony between distant brain regions (i.e. inter-hemispheric connectivity^14^ or fronto-parietal connectivity^15^) and that this modulation of inter-areal synchrony affects cognition. These studies applied tACS in such a way that two brain regions were stimulated with currents being either completely in-phase (i.e. 0 deg phase difference between region A and region B), or fully out-of-phase (180 deg phase difference between A and B; see Figure 2A). These results suggest that it is possible to control the degree of phase synchrony between brain regions via entrainment of oscillations with tACS. However, it should be noted that one study failed to show evidence for such entrainment effects and suggested that tACS rather affects oscillations via inducing changes in synaptic plasticity as opposed to entrainment^16^. Another important open question is whether with tACS it is possible to affect brain oscillations in a temporally sensitive, i.e. transient manner. Importantly, previous studies which demonstrated entrainment effects on behaviour mostly applied tACS in a sustained way over 10 – 30 minutes^17-19^. Therefore, it is unclear whether tACS is capable of targeting brain oscillatory activity in a temporally sensitive way which matches the temporal dynamics these oscillations naturally fluctuate in during cognitive tasks, which is in the range of fractions of seconds rather than minutes.

In the current study we addressed two issues. The first issue, building on our previous EEG-fMRI findings^4^, addresses the question of whether the manipulation of prestimulus interregional phase synchrony at 7 Hz between lower and higher order visual processing regions modulates perceptual integration. Perceptual integration refers to a process of transforming distributed activity in lower visual regions into meaningful object representations, by integrating neural information across object features^20^ or across space^21^. The process involves the bottom-up signalling of candidate features into a spatial map^22^ as well as the top-down selection of targets based on their spatial location^23^. Thus, perceptual integration relies on crosstalk between cortices on different levels of the visual processing hierarchy, i.e. regions in the occipital and parietal cortex.

The second issue concerns the question of whether tACS is capable of targeting oscillatory activity in a temporally specific way, i.e. whether it is possible to entrain oscillatory activity within a very short period of time (1 -1.8 seconds). To test these two hypotheses we applied tACS in five different ways (see Figure 2A). To this end, we either stimulated the left occipital and right parietal regions to be (i) perfectly in-phase, or (ii) out-of-phase (the stimulated regions were derived from a previous EEG-fMRI study^4^). Additionally, each region was stimulated separately (iii-iv). Finally, a sham stimulation condition (v) was carried out. We predicted that perceptual integration shows a strong phase modulation (i.e. tACS phase at stimulus onset) in the in-phase condition, as in this condition the windows for communication would – ideally – be perfectly “open” or “closed”, therefore the likelihood of whether a stimulus is perceived should strongly depend on the prestimulus phase. In contrast, in the out-of-phase condition no modulation of perception by prestimulus phase should be obtained, as the two phases cancel each other out. The two local stimulation conditions (i.e. right parietal, left occipital) might show a medium level of phase modulation. Finally, we also expected that synchronizing/de-synchronizing the two involved brain regions in the prestimulus interval should influence the general level of perceptual integration, irrespective of phase at stimulus onset, via biasing the two brain regions towards best or worst case scenarios for neural communication^24,25^ (see Figure 2C). In order to test for effects of entrainment we used two dependent variables, (i) behaviour and (ii) post stimulation EEG activity phase-locked to tACS offset (i.e. oscillatory entrainment echoes^26^).

**Figure 2.**
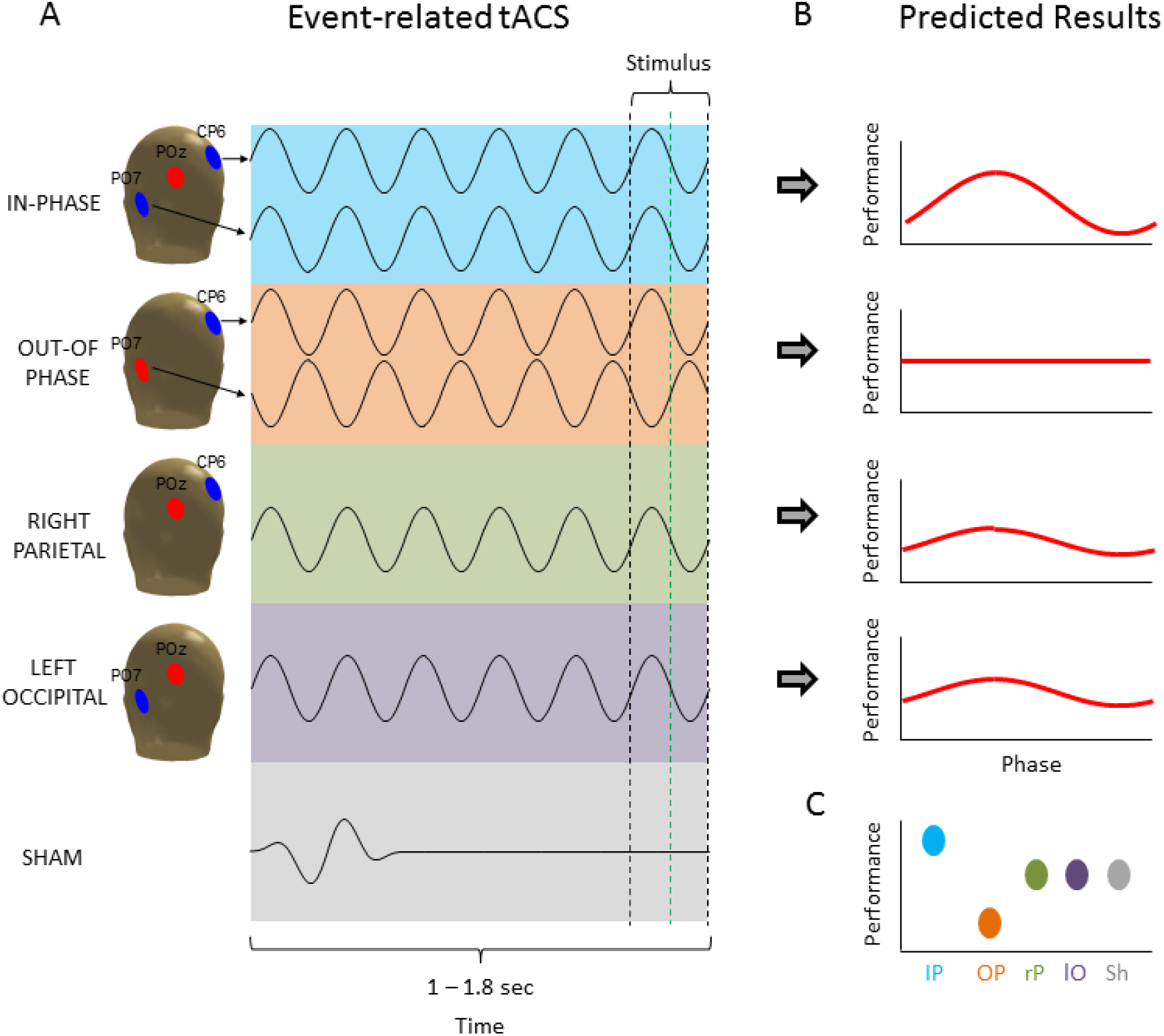
The five different tACS conditions are shown (A) together with the predicted results (B-C). (A) TACS was always delivered with a frequency of 7 Hz and an amplitude of 1 mA. The four active tACS conditions differed only in terms of which electrode locations were used for stimulation. Red electrodes positions indicate the reference (i.e. return) electrode whereas blue electrodes indicate the active (i.e. stimulation) electrode/s. For SHAM a short (~282 ms) 7 Hz sine wave was applied varying randomly between the four possible electrode combinations. The stimulus was presented at the end of stimulation at different phase angles. TACS was terminated at the earliest possible zero crossing occurring after stimulus onset (i.e. the green line for approx. half of the trials). This way the stimulus only overlapped for a maximal duration of 71 ms with tACS stimulation. (B) The predicted results in terms of phase modulation are shown. Strongest phase modulation was predicted for the IN-PHASE, weakest phase modulation was expected for the OUT-OF-PHASE modulation, with RIGHT-PARIETAL and LEFT-OCCIPITAL in between. (C) The predicted outcomes for general perception performance (i.e. independent of phase) are illustrated.

## Results

### General effects of tACS on perceptual integration

Across all conditions the average hit rate was 0.76 (s.d. 0.10) and the average correct rejection rate was 0.72 (s.d. 0.18). This shows that the pretesting session, where the difficulty of the task was adjusted for each participant to yield performance of around 0.75 was successful (see Methods). To assess the subjects ability to discriminate between targets and distractors, false alarms were subtracted from hits (Hits – FAs) and subjected to a repeated measurement ANOVA with the factor stimulation (IN-PHASE, OUT-OF-PHASE, RIGHT-PARIETAL, LEFT-OCCIPITAL, and SHAM). The sphericity assumption was not violated by our data (χ^2^(9) = 8.42, p = .49) therefore no correction for sphericity violation was applied. A marginally significant main effect was obtained (F_4,80_=2.48, p = 0.05). The results are shown in Figure 3A. Best performance was obtained for IN-PHASE (0.511), followed by OUT-OF-PHASE (0.498), RIGHT PARIETAL (0.492), SHAM (0.468) and LEFT OCCIPITAL (0.455). Post-hoc tests revealed that only performance in the IN-PHASE condition differed significantly from SHAM (t_20_=2.92; p<0.01; two-tailed), no other comparison reached significance (t’s<1.53; p’s>0.14). While the enhanced performance for IN-PHASE compared to SHAM is in line with our hypothesis, the almost equal performance in the OUT-OF-PHASE compared to the IN-PHASE condition (t_20_=0.62; p=0.54) is not. A post-hoc interpretation that would fit the pattern of results is that stimulation conditions that included the right parietal electrode (i.e. IN-PHASE, OUT-OF-PHASE, and RIGHT PARIETAL) lead to better performance compared to conditions that do not include this site or are ineffective (i.e. SHAM, LEFT OCCIPITAL). Indeed averaging performance in such a way revealed a significant difference between right parietal electrode stimulation sites (IN-PHASE, OUT-OF-PHASE, and RIGHT PARIETAL) compared to SHAM and LEFT OCCIPITAL (0.50 vs. 0.46, t_20_=2.69; p<0.05; two-tailed).

**Figure 3.**
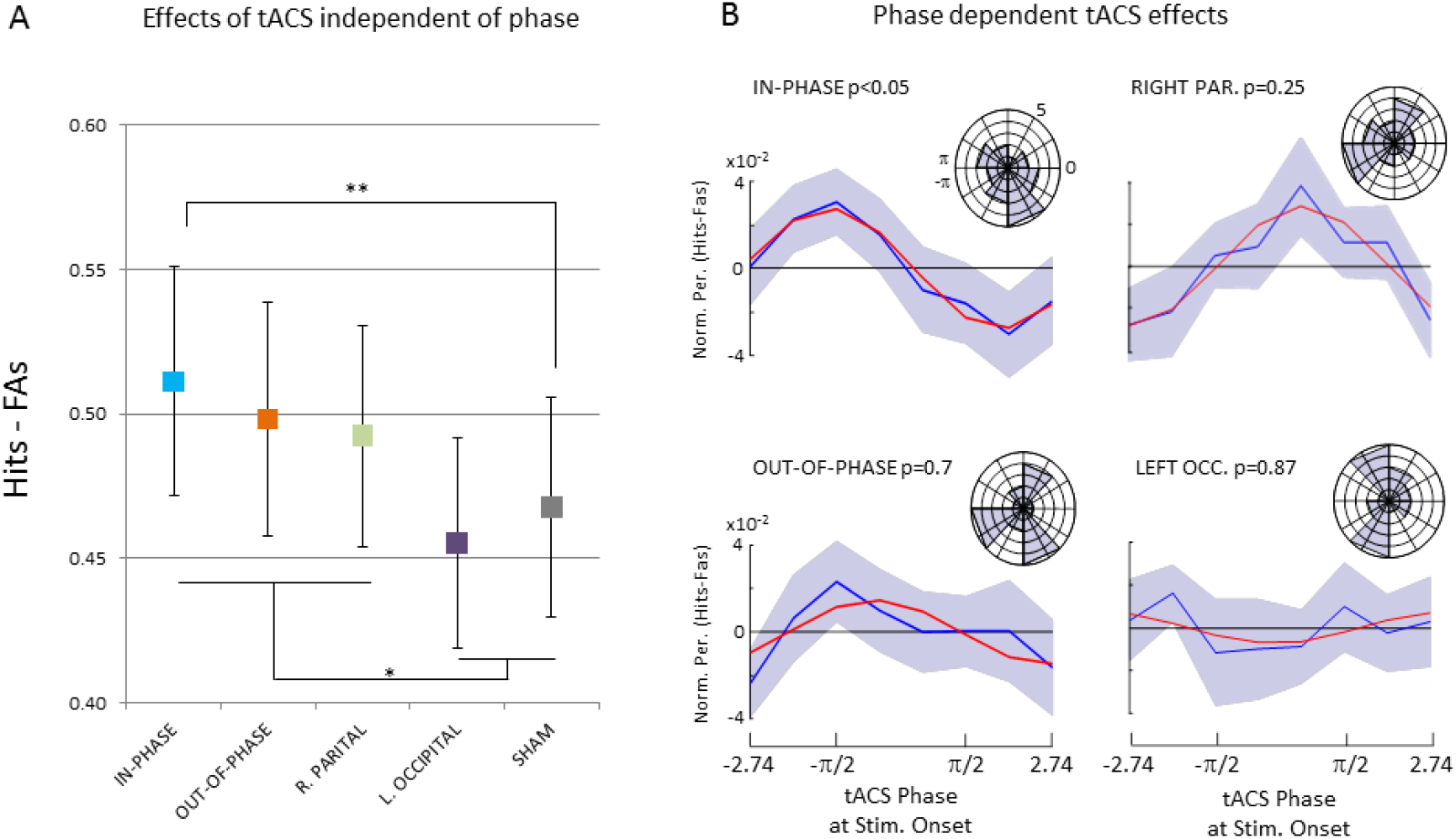
Behavioural results are shown. (A) Perceptual performance is shown as the difference between Hits minus False Alarms (FAs) for the five different stimulation conditions independent of phase. (B) Normalized (i.e. zero centred) and smoothed average perception performance is shown separately for the four active stimulation conditions across the 8 phase bins. P-values reflect the type-II error probabilities of that the data followed a circular pattern as obtained by non-parametric statistical testing (see Methods). Blue lines reflect the averaged data, red lines show the best fitting sine wave. The circular histograms plot the distribution of the “best phase” (i.e. phase bin with highest performance) across subjects. Error bars and shaded areas reflect mean standard errors. *p<0.05; ** p<0.01

### Phase dependent effects of tACS on perceptual integration

To test whether perceptual integration depended on the tACS phase at stimulus onset the behavioural data for each condition were sorted into 8 phase bins. Perceptual performance was again calculated as the difference between Hits and FAs (Hits - FAs). To reduce noise, a smoothing procedure was applied, using a running average with a window size of 3 bins (see Methods) and a sine wave was fitted to the averaged data across subjects for each of the four active stimulation conditions. The results are shown in Figure 3B. A non-parametric randomization test (see Methods) revealed that only perception in the IN-PHASE condition could be accounted for by a sine wave (p<0.05). None of the other conditions showed a significant sine wave fit on the averaged data (p’s>0.25). To further explore the relationship between phase and perception performance a Raleigh Test for non-uniformity was carried out on the “best fitting” individual phase, i.e. the phase that on an individual level corresponded to the best performance. However, no significant deviation from non-uniformity was obtained for any of the four conditions. Phase histograms are plotted in Figure 3B for descriptive purposes. Results for unsmoothed data are shown in Figure 4.

**Figure 4.**
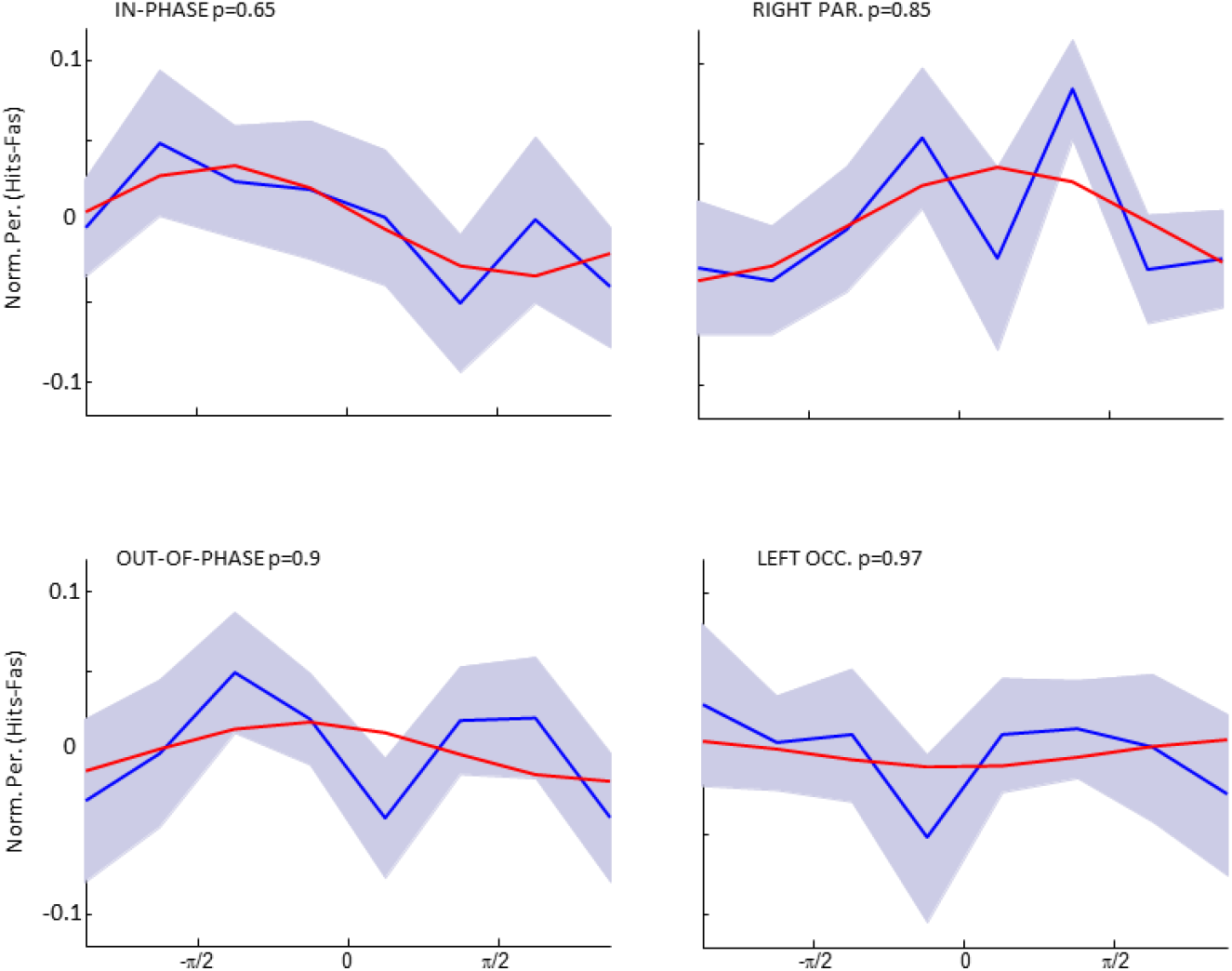
The unsmoothed averaged data for each stimulation condition is shown together with the best fitting sine wave and the p-level of the sine wave fit.

### TACS entrainment echoes

Possible aftereffects of tACS on EEG oscillatory activity were explored using the same logic as in one of our previous studies^26^. The main assumption being if the IN-PHASE tACS indeed entrained a theta oscillation we should see an after effect of that entrainment (“entrainment echo”) in the EEG phase-locked to the offset of tACS. To this end we calculated power spectra of the ERP locked to the offset of tACS and tested whether a stronger theta aftereffect was present for the IN-PHASE condition compared to OUT-OF-PHASE. We focused on these two conditions for two reasons: (i) according to our hypothesis, these conditions should show the strongest difference in theta phase-locking and (ii) they produced a comparable magnitude of EEG artefacts induced by tACS (i.e. SHAM wouldn’t be a good control condition, as tACS artefacts were substantially reduced). The results of this analysis are shown in Figure 5. A non-parametric randomization test (see Methods) revealed stronger evoked theta power (6-8 Hz) for the IN-PHASE compared to the OUT-OF-PHASE condition. This effect was significant in a parietal cluster of 9 electrodes (p_corr_<0.05), and showed a non-significant trend for a cluster of 3 fronto-central electrodes (p_corr_=0.11). This parieto-fronto-central topography is in line with the topography of the prestimulus phase effect observed in our previous study^26^. None of the other tested frequencies (2-15 Hz) revealed a significant difference. Unexpectedly, the maximum difference occurred 1 Hz above the actual stimulation frequency (i.e. 8 Hz compared to 7 Hz).

**Figure 5.**
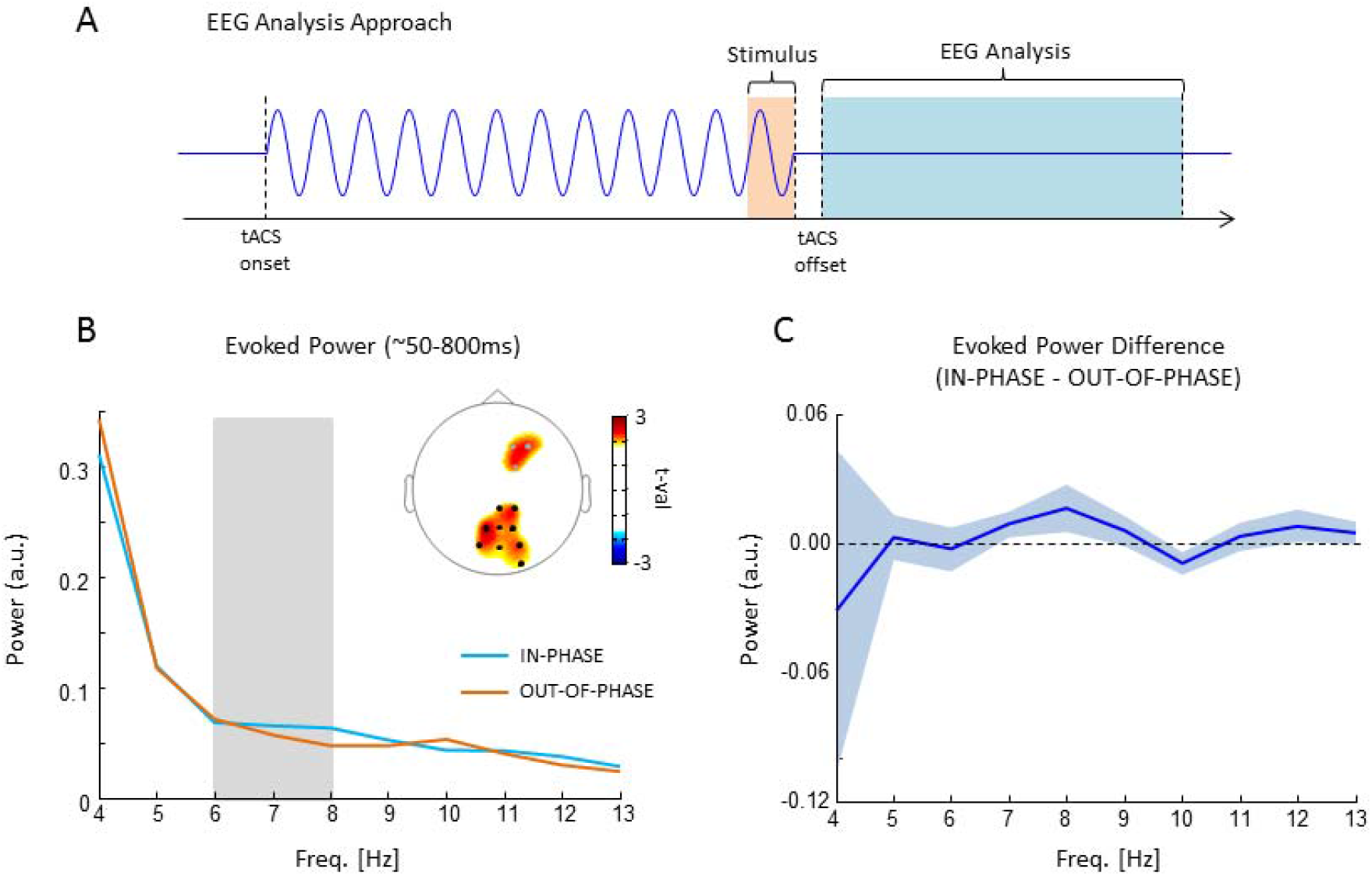
EEG results are shown. (A) The analysis approach for EEG is illustrated. EEG epochs were time locked to tACS offset (as opposed to stimulus onset). The time window varied slightly from subject to subject depending on the artefacts introduced by tACS but on average corresponded to length of 880 ms starting ~90 ms after tACS offset (light blue). The exact time windows were adjusted individually due to variable delays in tACS offset artefacts (see Methods). Evoked power was calculated as the power spectrum of the resulting ERP. (B) Evoked power is shown for the IN-PHASE and OUT-OF-PHASE condition. The topography shows significant differences (black dots = p_corr_<0.05; grey dots = p_corr_=0.11). (C) The difference between IN-PHASE and OUT-OF-PHASE in evoked power is shown averaged across the sign. parietal electrodes (i.e. black dots in B). Shades areas represents mean s.e.

## Discussion

We here employed a new tACS protocol in theta frequency range to address two questions: (i) Is the role of prestimulus inter-areal phase synchrony causally relevant for perceptual integration? (ii) Can tACS be used to entrain oscillations in a temporally sensitive manner in order to target oscillatory dynamics at a time resolution that matches cognitive time scales? Since we feel that our data does not warrant a clear answer to either of those questions, the aim here is to discuss the results in a balanced manner such that supportive and counter-supportive arguments are presented. We hope that this will trigger of follow-up experiments which further clarify the above questions. In the following, we will first discuss the effects of tACS on behaviour, i.e. general performance and phase modulation, and then the effects on the EEG (i.e. entrainment echoes), followed by a consideration of important limitations.

Concerning the general effect of tACS on perceptual integration, we predicted a difference between the five stimulation conditions, such that performance would be highest for the synchronizing (in-phase) stimulation and lowest for the de-synchronizing stimulation (out-of-phase), with the other conditions being in between (see Figure 2c). Indeed, our results did show a significant main effect of stimulation on perceptual integration, and in-phase stimulation showed the highest levels of performance. However, out-of-phase stimulation quite clearly did not impair perception performance. On a post hoc level these results are more in line with the interpretation that stimulation montages which include the right parietal cortex increase performance. This idea would be consistent with the results obtained in our previous EEG-fMRI study^4^, which implicated the right intra-parietal region in this task. Such a post-hoc interpretation is also consistent with several other studies, indicating the right intraparietal sulcus to be a critical region for perceptual integration^20,27^. However, this post-hoc interpretation remains to be tested by future studies. Coming back to the two main questions, these first results do not support the idea that prestimulus interareal theta phase is causally relevant for perceptual integration, as in-phase and out-of-phase did not show any difference in performance. However, the fact that short lived prestimulus tACS did modulate perceptual performance suggests that tACS can be used in a time sensitive manner. Although the exact nature of the effect of prestimulus theta stimulation on perceptual integration is unclear this is an interesting avenue for future tACS experiments.

Central to the hypothesis that interareal prestimulus theta synchronization is causally relevant for perceptual integration are the phase modulation results shown in Figure 3B. In line with our hypothesis the in-phase stimulation condition did show a significant phase modulation of perception performance, whereas none of the other stimulation conditions did. Thus, theta phase at stimulus onset may be causally relevant for perception and may be modifiable by transient tACS. Such a result would indeed be exciting news for the tACS field as it suggests a new way of using tACS to modify brain oscillations on a time scale that matches with cognitive operations. However, several aspects in our data warrant caution to not over interpret these results. First, the phase modulation in the in-phase stimulation condition was only obtained after smoothing the data with a running average. Although it is not uncommon in the field to apply such smoothing procedures^28^, and although we did take care that the randomization statistics were not biased by smoothing itself, it is an important aspect to consider. Furthermore, we did not find evidence for a clustering of the “best” phase (i.e. phase with highest performance) across participants in the in-phase condition which further calls into question whether the synchronizing theta tACS did indeed modulate perception performance in a phase dependent manner. Finally, several attempts to directly test for differences of the magnitude of phase modulation (i.e. goodness of sine wave fit, ANOVAs, etc.) between the different stimulation conditions failed, which also violates our predictions. Together, although there are some aspects in the behavioural data that are in line with prestimulus theta tACS entraining perception, several aspects do not support this conclusion. This might lie in the fact that tACS effects on behaviour are quite subtle due to tACS being a rather weak stimulation technique, especially when applied only for a very brief time. This necessitates estimations of effect sizes in order to determine the appropriate sample sizes and number of trials before running similar tACS studies in the future, for which the data reported here should be helpful.

Importantly, in the current study we did not only measure behaviour but also recorded EEG aftereffects in order to test for effects of prestimulus tACS. Specifically, we tested whether the EEG shows entrainment echoes, which are short-lived oscillatory aftereffects that are phase locked to the offset of rhythmical stimulation (see^26^ for an example of entrainment echoes in the beta frequency range in the prefrontal cortex). If prestimulus tACS does indeed synchronize or de-synchronize large scale neural ensembles in the theta frequency range then stronger ERP power should be observed in the entrained frequency for the in-phase compared to the out-of-phase stimulation condition. This hypothesis is supported by our EEG results (see Figure 5). The observed topography is consistent with the site of stimulation (i.e. parieto-occipital areas) and is also consistent with the topography of the prestimulus theta effect reported in our previous study that used exactly the same task^4^. This observed tACS aftereffect in ERP power was frequency specific and consistent with the stimulated frequency, albeit 1Hz higher. We are therefore inclined to interpret this effect as an entrainment echo, i.e. short-lived oscillatory signal in the entrained frequency that is phase-locked to tACS offset. However, some important limitations need to be considered. During tACS the EEG signal was massively contaminated by tACS and saturated on several electrodes. After tACS offset the EEG drifted back to baseline, which induced a strong artefact in the lower frequencies (see Figure 5B). A second limitation is that our task design only left a small window (880 ms) for analysis of tACS aftereffects, which precluded a more sensitive time-frequency analyses in the 7 Hz frequency (due to temporal smearing). Although these limitations do not necessarily render the observed frequency specific entrainment echo spurious, future studies should take these issues into account. Together, the EEG results seem to be the strongest evidence reported here in favour for the hypothesis that tACS can entrain neural oscillations in a temporally sensitive manner. This latter finding contradicts a previous tACS-EEG study where alpha oscillations were stimulated and no entrainment echoes were observed^16^. The reasons for this discrepancy might lie in the several differences between the two studies such as placement of the stimulation electrodes, differences in the stimulation protocol (i.e. intensity, size of electrodes, etc.), cognitive task (i.e. perception task vs. rest), or frequency of stimulation (i.e. 7 Hz vs. 10 Hz).

Finally, an important issue to consider when utilizing tACS protocols which synchronize or de-synchronize distant brain regions is that such stimulation could not only bias two regions (i.e. frontal and parietal^15^) to exhibit a different phase, but might in fact drive completely different brain regions. In the present study, the out-of-phase condition involved current being passed between left occipital and right parietal regions (see figure 2). We cannot rule out the possibility that in this stimulation condition different brain areas are being stimulated, i.e. the tissue between the active and the return electrode, compared to the in-phase condition. Although modelling studies suggest that, during transcranial direct current stimulation, the electrical field is strongest close to the anode^29^, the distance and location of the cathode has to be taken into account as well^30-32^. As the electrical field strength is strongest close to the electrode edges between the anode and the cathode, the different relative location of the return electrode in the out-of-phase condition as compared to the in-phase condition could have resulted in a different distribution of the electrical field. Unfortunately, our knowledge about the current distribution applied by different electrode montages is rather scarce, because the modelling of currents is quite complex^33^ and at the moment these models have not been validated by experimental data.

## Summary

We here presented a study where we applied transient tACS in the prestimulus interval during a perceptual integration task in order to test for a causal involvement of theta interareal phase synchrony for perception. On a more general level our study also tested whether short lived tACS has an effect on behaviour and EEG oscillations. Together, the behavioural results do not strongly support the hypothesis that prestimulus theta phase plays a causal role for perceptual integration. However, our results do suggest that transient tACS is capable of inducing entrainment echoes and has an effect on behaviour.

## Methods

### Subjects

24 subjects participated in the experiment (mean age 24.36 ± 5.1, 12 males). Three subjects were left handed. The data of three participants were discarded from further analysis due to strong EEG artifacts, leaving a sample of 21 participants for data analysis (mean age 24.1 ± 5.3, 10 males). All participants were screened for possible risk factors of transcranial current stimulation (TCS)^34^, had normal or corrected to-normal vision and reported no history of neurological disease or brain injury. Subjects were paid £21 or received course credits for participation. The study was approved by the ethics committee of the University of Birmingham.

### Stimuli

To probe the effects of synchronizing and desynchronizing tACS on visual perception a similar perceptual integration task as in our previous study was used^4^. Stimuli consisted of arrays of Gabor elements, subtending 14 by 14 degrees of visual angle, that did or did not contain a path of collinear-oriented elements (’’contour’’ or ‘’non-contour’’; see Figure 1). The stimuli were generated with a procedure similar to that of Watt et al^35^.

In a first step, a path with nine line segments was constructed for supporting the later placement of the contour elements. The first segment had a random orientation, and each following segment was rotated by adding or subtracting a constant value to the angle of the previous line segment. This value, the path angle, was adjusted individually for each subject (details see below). If a path contained a loop, it was discarded and a new solution was computed. One Gabor element was placed in the middle of each line segment, oriented collinear to the path. The distance between neighbouring elements was 1.2 ± 0.18 degrees of visual angle. The contour was copied to a random location within the stimulus array, and then randomly oriented Gabors were added to the array until no more elements could be placed. Constraints were that the distance between the Gabors was at least 1 degree of visual angle from center to center, and that no Gabor fell in the screen center where a fixation mark would occur. The resulting displays contained ~130 Gabors on average. All Gabor elements had a spatial frequency of 2.4 cycles per degree. The Michelson contrast was set to 0.9, and the standard deviation of the Gaussian envelope, controlling the size of the Gabor, was 0.12 degrees of visual angle.

For constructing non-contour displays, we used the same algorithm as for the construction of contour displays but rotated adjoining Gabor elements by plus or minus 45 degrees. Thus, non-contour displays resembled contour displays with respect to spacing, positioning and the number of elements, but did not contain a contour path.

### Procedure

Each participant participated in two sessions: a training session and the main experiment. For both sessions the same experimental setup was used. Participants were seated at 80 cm distance from a 19 inch monitor (resolution: 1280x1024 pixels, 60 Hz frame rate). Stimuli were presented in a grey background on the center of the screen using the Psychophysics Toolbox extension for Matlab^36^. In each session Gabor arrays were presented for 200ms and subjects were given 1 second to respond via a button press on a computer keyboard whether they did perceive a contour or not. Responses were given with the right index and middle fingers. In both sessions 50% of the stimuli contained a contour stimulus. After the response period, a fixation-cross appeared for a variable and unpredictable duration between 1000 and 1800 ms. In the main experiment this fixation cross period was the period that was used for tACS.

### Training session

The training session was performed one day before the main experiment. The main purpose of the training session was to adjust the difficulty of the task individually such that each participant reached a performance close to 75% accuracy. Difficulty of the task was adjusted via varying the variation in path angle between contour forming elements, i.e. if this variation is small a contour can easily be detected and detection gets more difficult with increasing variation. The training was started with low difficulty trails (small variation in path angle) and increasing difficulty after every block (i.e. 40 trials) of ≥75% accuracy. The training session was stopped after 3 blocks in a row of stable ≤75% accuracy. However, lowest number of blocks that participants had to finish was 10. The resulting path angle was then used to create stimuli for the respective participant for the main experiment.

### Main experiment

The same trial structure was used for the main experiment as for the training session. Four blocks of 200 trials were carried out with short breaks in between to make sure the participants are feeling ok and are awake. Half of the trials contained a contour, whereas the other half did not. Overall 800 stimuli were presented overall during the main experiment which lasted ~ 45 minutes.

### TACS parameters

Transcranial Alternating Current Stimulation (tACS) was delivered via a 4 channel DC Stimulator MC (NeuroConn, www.neuroconn.de). Stimulation was synchronized to the task such that tACS was only delivered during the interstimulus interval (see Figure 1). In all conditions a 7 Hz sine wave was used for stimulation at an intensity of 1mA (i.e. 2 mA peak to peak). Stimulation was applied via round rubber electrodes with a diameter of 3.7 cm (10.75 cm^2^, NeuroConn, Ilmenau, Germany) resulting in a current density of 0.093 mA/cm^2^. Stimulation electrodes were centered at the following three EEG electrode positions: PO7, POz and CP6 (see Figure 6). These positions were derived from a neuro-targeting software (Soterix Medical Inc, New York, USA) which uses a finite-element model of an adult male brain to model the current distribution in the brain. Stimulation sites were chosen to result in the highest target field intensity in left middle occipital cortex (MNI coordinates: x=-32, y=-89, z=18; BA 19) and right inferior parietal lobule (MNI coordinates: x=44, y=-64, z=44; BA 40), guided by the results of our previous EEG-fMRI study^4^. Another criterion for choosing the stimulation sites was that the same return electrode could be used for both regions, to allow for “zero-phase” and “180-phase” lag stimulation^15^ (see below). Conductance between scalp and electrode was established via Ten20^®^ paste and resistance was kept below 5kohm. Before the start of the main experiment participants were familiarized with tACS and desensitized to the stimulation intensity to avoid adverse reactions. To this end, trains of 2000 ms 7Hz tACS were delivered starting at an intensity of 0.4 mA and increasing in steps of 0.2 mA to the resulting stimulation intensity of 1 mA. This adaption procedure lasted for ~10 minutes and most participants reported little to no sensation during stimulation after this adaptation phase.

**Figure 6.**
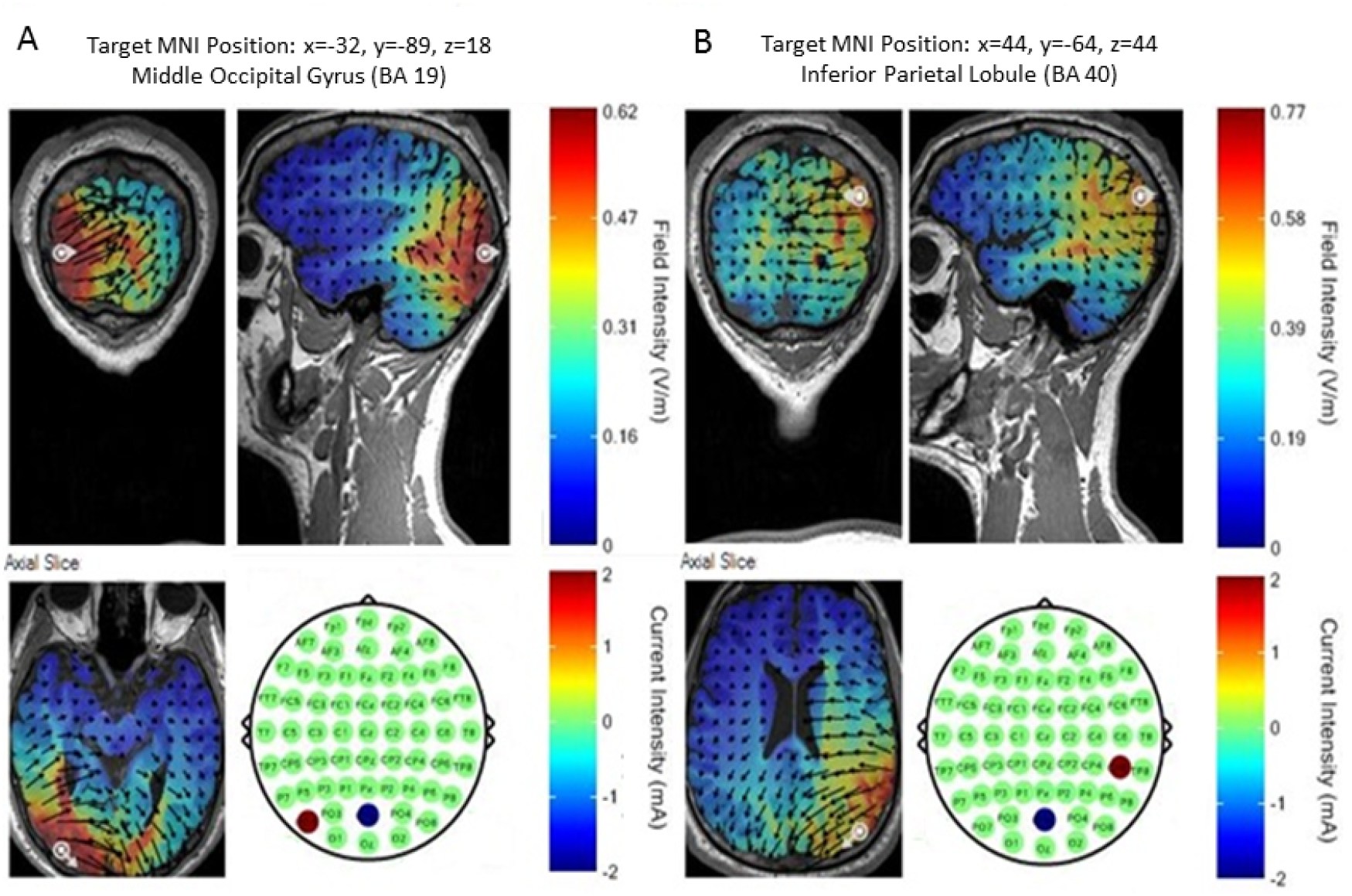
Stimulation electrode configurations are shown for left middle occipital gyrus (A) and right inferior parietal lobule (B). Current field intensity is shown using a finite element model as provided in the software SOTERIX^®^.

Five conditions of tACS were conducted which differed in terms of the site of stimulation, phase lag between stimulation sites and duration (see Figure 2A). The five stimulation conditions were: IN-PHASE, OUT-OF-PHASE, RIGHT PARIETAL, LEFT OCCIPITAL, and SHAM. IN-PHASE tACS was implemented by simultaneously alternating electrical fields between the stimulation electrodes placed at PO7 and CP6, using POz as a “return” for both electrodes, thus resulting in a 0 degree phase shift between these two alternating fields^15^ (see Fig. 2A). OUT-OF-PHASE tACS was implemented by alternating electrical fields between the electrodes placed at PO7 and Cp6 such that the phase between the two sites (and presumably in the underlying regions) showed a 180 degree phase difference at any point in time^15^ (see Fig. 2B). These two stimulation conditions were of main interest for our experiment. To control for unspecific effects of tACS (i.e. phosphenes, tingling sensation, etc.) two active control conditions were performed, where either one of the two regions were stimulated, i.e. RIGHT PARIETAL (Fig. 2C), or LEFT OCCIPITAL (Fig. 2D) at the same intensity and same duration as in the IN-PHASE and OUT-OF-PHASE conditions. Lastly, a SHAM condition was carried out which consisted of one 7 Hz cycle ramping up and another cycle ramping down, resulting in ~286 ms of stimulation at the beginning of fixation period (Fig. 2E). Sham stimulation was randomly performed for all four electrode montages (i.e. IN-PHASE, OUT-OF-PHASE, RIGHT PARIETAL, and LEFT OCCIPITAL). Conditions varied randomly on a trial-by-trial basis with a different random sequence generated for each subject.

Of crucial interest was the question whether perception performance depended on the phase at the onset of the visual stimulus. It was therefore crucial to control for the phase when the stimulus was presented on the screen. To this end, we factored in the timing delays and jitters of our stimulation setup and aimed at adjusting our protocol such that the phase at stimulus onset was uniformly distributed at stimulus onset. However, due to unpredictable jitters (+/- 17 ms) with respect to stimulus onset it was necessary to back-sort the trials after the experiment according to the phase at stimulus onset, covering 8 equally sized phase bins centred at 202.5°, 247.5°, 292.5°, 337.5°, 22.5°, 67.5°, 112.5°, 157.5°. Thus, out of the 800 trials, 160 trials were available for each of the five tACS conditions (IN-PHASE, OUT-OF-PHASE, …, SHAM), 20 trials were available for each phase bin (202.5°, …, 157.5°), of which half contained contour stimuli and half contained non-contour stimuli. For phase estimation at stimulus onset the current waveform of the stimulator output was used. Phase was calculated by a Hilbert transform. After the experiment participants were asked to evaluate tACS side effects (how painful the stimulation was, intensity of phosphenes and other visual artifacts) in a such scale: 0 – No, 1 - Mild, 2 – Moderate, 3 - Strong, 4 - Severe, 5 – Unbearable. For the ratings of phosphenes and other visual artifacts only 13 datapoints were available, because these ratings were introduced at a later point. Averaged ratings and statistical analysis of the ratings are reported in Supplementary Materials.

### Questionnaire on adverse effects

To obtain possible side effects of the stimulation protocol, an adapted version of the questionnaire proposed by Brunoni et al^37^ was used. The following side effects were assessed: headache, neck pain, itching, sleepiness, trouble concentrating, acute mood change, fatigue, nausea, muscle twitches in face or neck, tingling sensation in head or on scalp, phosphenes, burning sensation in head or on scalp, epileptic seizure, non-specific uncomfortable feeling. Participants rated if they experienced these side effects and if yes, how strong they experienced them (mild, moderate, severe). The most common side effects reported were tingling sensation (85.7%), sleepiness (71.4%), trouble concentrating (66.7%) and fatigue (61.9%). Participants also experienced non-specific uncomfortable feeling (42.9%), burning sensation (38.1%), itching (38.1%), phosphenes (23.8%), headache (19.1%) and neck pain (14.3%). Individual subjects reported acute mood change, nausea and muscle twitches. Overall 10 subjects indicated that one or more of these effects might have had an impact on their performance.

### EEG recording

The EEG was recorded throughout the experiment to test for after effects of tACS after stimulation offset (i.e. entrainment echoes^26^). The EEG was recorded using a 64 channel NEURO-PRAX^®^ Amplifier (NeuroConn, Ilmenau, Germany). 61 Ag/Cl Electrodes were placed on the scalp according to the international 10-20 system mounted into an elastic cap. EEG signals were sampled at 1000 Hz and referenced to the FP1 electrode, because this electrode showed the lowest artifacts from tACS. Impedance was kept at optimal values as suggested by color coding of the EEG recording system (discrete resistance values kohm were not accessible). A high-chloride, abrasive electrolyte-gel was used (Abralyt HiCl, Easycap^®^, Herrsching, Germany).

### Behavioral Data Analysis

Behavioral responses were classified into four categories: Hits (H), i.e. correctly identified contours; Misses (M), i.e. contour stimuli incorrectly endorsed as non-contour stimuli; Correct Rejections (CR), i.e. correctly identified non-contour stimuli, and False Alarms (FA), i.e. non-contour stimuli incorrectly endorsed as contour stimuli. Perceptual performance (P) was quantified as the ratio of Hits minus the ratio of False Alarms according to the following equation:

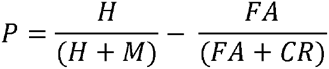

This measure quantifies the degree to which participants were able to discriminate between contour and non-contour stimuli.

In a first step, P was calculated for each of the five tACS conditions, irrespective of phase at stimulus onset, and compared using a one factorial Repeated Measurement ANOVA to test for global differences between stimulation conditions. Sphericity was tested using Mauchly’s test of Sphericity. Post-hoc comparisons were conducted using Paired-Sample T-Tests. In a second step, and of main interest for our hypotheses, P was calculated for every phase bin in every active tACS condition (i.e. IN-PHASE, OUT-OF-PHASE, RIGHT PARIETAL, and LEFT OCCIPITAL). No phase binning was carried out for the SHAM stimulation condition. The phase binned data for each condition and participant was zero centered by subtracting the mean across phase bins. Furthermore, the behavioral data was smoothed using a moving average with a window size of 3 bins, such that P for a particular phase bin, i.e. 22.5° = 337.5° + 22.5° + 67.5° / 3. This smoothing procedure has two effects on the data of which one is desirable and the other is problematic. On the upside smoothing can reduce noise in the data and benefit cases where a low trial number per bin is present (as we argue is the case in our data). However, on the downside smoothing introduces biases, i.e. dependencies between neighbouring data points, and can render non-circular data (i.e. were a local maxima is present) circular. Hence, smoothing biases the data to find circular patterns which is an important caveat to consider. We also carried out all calculations on the unsmoothed data and found a similar pattern of results but none of which survived statistical testing (see below).

In order to statistically quantify whether perception performance was modulated by the phase of tACS at stimulus onset we fitted a sine wave on the smoothed phase binned data averaged across participants, using the curve fitting toolbox in MATLAB, and the explained variance (R^2^) value was retained. Therefore we assumed a consistent relationship across participants between phase and perception, following our previous results from EEG-fMRI^4^. To test this sine wave fit against the null hypothesis (i.e. performance does not follow a circular pattern) we carried out a non-parametric randomization test. To this end the unsmoothed but zero-centered data was shuffled randomly across phase bins and then smoothed. It is crucial to carry out smoothing after the data was shuffled as otherwise the randomization statistics would be heavily biased towards rejecting the null-hypothesis. This procedure alleviates to some extent the smoothing problematic mentioned above. Finally, a sine wave was fitted on the average data across participants, and the explained variance value (R^2^) was retained. Parameters for smoothing and sine wave fitting were the same as for the real data. This randomization procedure was run 1000 times to create a distribution of R^2^ values under the null-hypothesis against which the R^2^ value from the real data was compared. Significance was assumed if the real R2 value was above the 95^th^ percentile of this distribution.

### EEG Data Analysis

EEG analysis was carried out using FieldTrip^38^ (http://fieldtrip.fcdonders.nl/) and in-house MATLAB scripts. Because the EEG data was heavily contaminated by tACS artifacts several preprocessing steps were required. The EEG was first resampled to 500 Hz and segmented into 1.2 second long trials starting 0.02 seconds before tACS offset. Then the EEG data was merged with the data from the tACS stimulation device to obtain the stimulation information for each trials (i.e. tACS condition and phase offset) and cut into shorted 960 ms epochs starting 40ms after tACS offset (tACS offset was accompanied by large drifts back to baseline). Only trials where a contour was presented and which resulted in a correct behavioral response were selected (i.e. Hits). The EEG data was then filtered between 0.5 Hz and 35 Hz using a FIR filter and polynomial trends were removed to reduce the slow drifts after tACS offset. After filtering the data was further cut such that 50ms at the beginning and end of the trial were removed. The exact time varied slightly between subjects depending on the artifacts and resulted in an average epoch length of 880 ms (s.d. 21 ms). The resulting data was first visually inspected and noisy EEG channels, as well as “dead” channels (i.e. channels that were located at stimulation electrodes) were rejected. Thereafter, the EEG data was re-referenced to Cz and subjected to an independent component analysis (ICA). Components reflecting eye movements as well as tACS artefacts (i.e. slow drifts surrounding the stimulation electrodes) were rejected. A further visual rejection was carried out and then the data was re-referenced to average reference and missing electrode positions were interpolated. Trials were then grouped into the two tACS conditions of interest, i.e. IN-PHASE and OUT-OF-PHASE. For each condition the trials were further grouped based on the phase of stimulation offset, i.e. whether the stimulation stopped at 0 degrees (zero crossing after trough) or 180 degrees (zero crossing after peak). This step was necessary to ensure that the phases do not cancel each other out when analyzing activity phase-locked to stimulation offset. To analyze possible after oscillatory aftereffects, i.e. entrainment echoes, time series were averaged for separately for the two tACS and phase offset conditions resulting in ERPs which were time-locked to tACS stimulation offset. These ERPs were then subjected to a fast fourier transform to obtain a frequency representation of the ERP signal. This analysis approach is the same that we used in a previous rTMS-EEG study to analyze entrainment echoes in the prefrontal cortex^26^. According to our stimulation frequency (7 Hz) our frequency region of interest was 6-8 Hz and differences between IN-PHASE and OUT-OF-PHASE were subjected to paired samples T-Tests. To control for multiple testing across electrode sites a non-parametric cluster permutation approach as implemented in FieldTrip (Monte Carlo method; 1,000 iterations; p<0.05) was used^39^. The observed clusters were considered significant when the sum of T-values exceeded 95% of the permutated distribution.

